# Restore mitophagy is essential to prevent cardiac oxidative stress during hypertrophy

**DOI:** 10.1101/2021.01.12.426366

**Authors:** Victoriane Peugnet, Maggy Chwastyniak, Steve Lancel, Laurent Bultot, Natacha Fourny, Olivia Beseme, Anne Loyens, Wilfried Heyse, Philippe Amouyel, Luc Bertrand, Florence Pinet, Emilie Dubois-Deruy

**Author notes:** **Corresponding authors:** Dr Emilie Dubois-Deruy and Dr Florence Pinet, INSERM U1167-IPL, 1 rue du professeur Calmette, 59019 Lille cedex, France. /.

## Abstract

Heart failure, mostly associated with cardiac hypertrophy, is still a major cause of illness and death. Oxidative stress causes contractile failure and the accumulation of reactive oxygen species leads to mitochondrial dysfunction, associated with aging and heart failure, suggesting that mitochondria-targeted therapies could be effective in this context. The purpose of this work was to characterize how mitochondrial oxidative stress is involved in cardiac hypertrophy development and to determine if mitochondria-targeted therapies could improve cardiac phenotypes. We used neonatal and adult rat cardiomyocytes (NCMs and ACMs) hypertrophied by isoproterenol (Iso) to induce an increase of mitochondrial superoxide anion. Superoxide dismutase 2 activity and mitochondrial biogenesis were significantly decreased after 24h of Iso treatment. To counteract the mitochondrial oxidative stress induced by hypertrophy, we evaluated the impact of two different anti-oxidants, mitoquinone (MitoQ) and EUK 134. Both significantly decreased mitochondrial superoxide anion and hypertrophy in hypertrophied NCMs and ACMs. Conversely to EUK 134 which preserved cell functions, MitoQ impaired mitochondrial function by decreasing maximal mitochondrial respiration, mitochondrial membrane potential and mitophagy (particularly Parkin expression) and altering mitochondrial structure. The same decrease of Parkin was found in human cardiomyocytes but not in fibroblasts suggesting a cell specificity deleterious effect of MitoQ. Our data showed the importance of mitochondrial oxidative stress in the development of cardiomyocyte hypertrophy. Interestingly, we observed that targeting mitochondria by an anti-oxidant (MitoQ) impaired metabolism specifically in cardiomyocytes. Conversely, the SOD mimic (EUK 134) decreased both oxidative stress and cardiomyocyte hypertrophy and restored impaired cardiomyocyte metabolism and mitochondrial biogenesis.

## Introduction

Heart failure (HF) remains a major cause of illness and death and its prevalence is increasing with a high rate of morbidity and mortality (Benjamin et al. 2017). Despite major significant advances, HF remains a therapeutic challenge, and several adverse consequences of HF are still poorly controlled. The common phenotype associated with HF is the development of cardiac hypertrophy, defined as an increase in heart size in order to compensate the increase in cardiac workload.

Oxidative stress, characterized by imbalanced reactive oxygen species (ROS) production and anti-oxidant defences, plays an important role in regulating a wide variety of cellular functions, including gene expression, cell growth and death (Tsutsui, Kinugawa, and Matsushima 2008). ROS cause contractile failure and structural damage in the myocardium (Tsutsui, Kinugawa, and Matsushima 2008) and activate a broad variety of prohypertrophy signalling kinases and transcription factors (Rababa’h et al. 2018). ROS production is well described to increase in several animal models of cardiac diseases (Emilie Dubois-Deruy et al. 2020) such as cardiac alterations associated with obesity (Jiménez-González et al. 2020), myocardial infarction (MI) (Merabet et al. 2011) or cardiomyocytes hypertrophy (Dai et al. 2011). Moreover, the accumulation of ROS in cells and tissues leads to mitochondrial dysfunction, defined as decreased mitochondrial biogenesis, mitochondria’s number and altered membrane potential (Bhatti, Bhatti, and Reddy 2017). Cardiac stress-induced mitophagy, the mitochondria selectively targeted autophagy, helps to remove damaged and dysfunctional mitochondria, thus preventing oxidative damage that could in turn initiate apoptosis and ultimately lead to HF (Morales et al. 2020). Indeed, impaired mitochondrial function is associated with aging and HF (Pisano et al. 2016), suggesting that mitochondria-targeted therapies could be effective in HF (Sabbah 2016; Senoner and Dichtl 2019).

In this context, mitoquinone (MitoQ) (Kim et al. 2020; Jiménez-González et al. 2020), a derivative of coenzyme Q and EUK 134 (Purushothaman and Nair 2016), an antioxidant with superoxide dismutase activity, have been demonstrated to effectively improve mitochondrial function and attenuate redox-related cardiomyopathies. Nevertheless, some studies, notably in cancer cells, described that MitoQ could lead to ROS production, rapid membrane depolarization and apoptotic cell death (Doughan and Dikalov 2007; Pokrzywinski et al. 2016; Gottwald et al. 2018). In this context, the purpose of this work was (1) to characterize how mitochondrial oxidative stress is involved in cardiac hypertrophy and (2) to determine if mitochondria-targeted therapies could improve cardiac phenotypes.

## Results

### Characterization of mitochondrial oxidative stress and biogenesis in hypertrophied neonatal rat cardiomyocytes (NCMs)

To understand the impact of oxidative stress in cardiac hypertrophy specifically in the cardiomyocytes, we used the model of hypertrophied NCMs (Turkieh et al. 2018). First, the cell index quantified by Real Time Cell Analysis (RTCA) technology was used to determine the time point from which the NCMs profiles were stable. Indeed, the curve shows the dynamic change in the cell index, which represents a relative change in electrical impedance depending on the proliferation of the cultured cells (first exponential phase) (Figure S1A). After serum privation, the curve stabilized and then we observed a decrease in cell index in early time of treatment of cardiomyocytes (120 min). Finally, the global cell index was similar between control (PBS) and hypertrophied NCMs (Isoproterenol (Iso)) (Figure S1A). Based on this data, we have chosen 24h of Iso treatment for the following experiments. We validated the development of hypertrophy in NCMs with a significant increase of cell area (Figure 1A) and observed a significant increase in mitochondrial superoxide anion levels quantified after 24h of hypertrophy with MitoSOX probe (Figure 1B) as well as a significant decrease in NADPH oxidase 4 (NOX4) (Figure 1C) without any mitochondrial hydrogen peroxide accumulation (Figure S1B).

**Figure 1:**
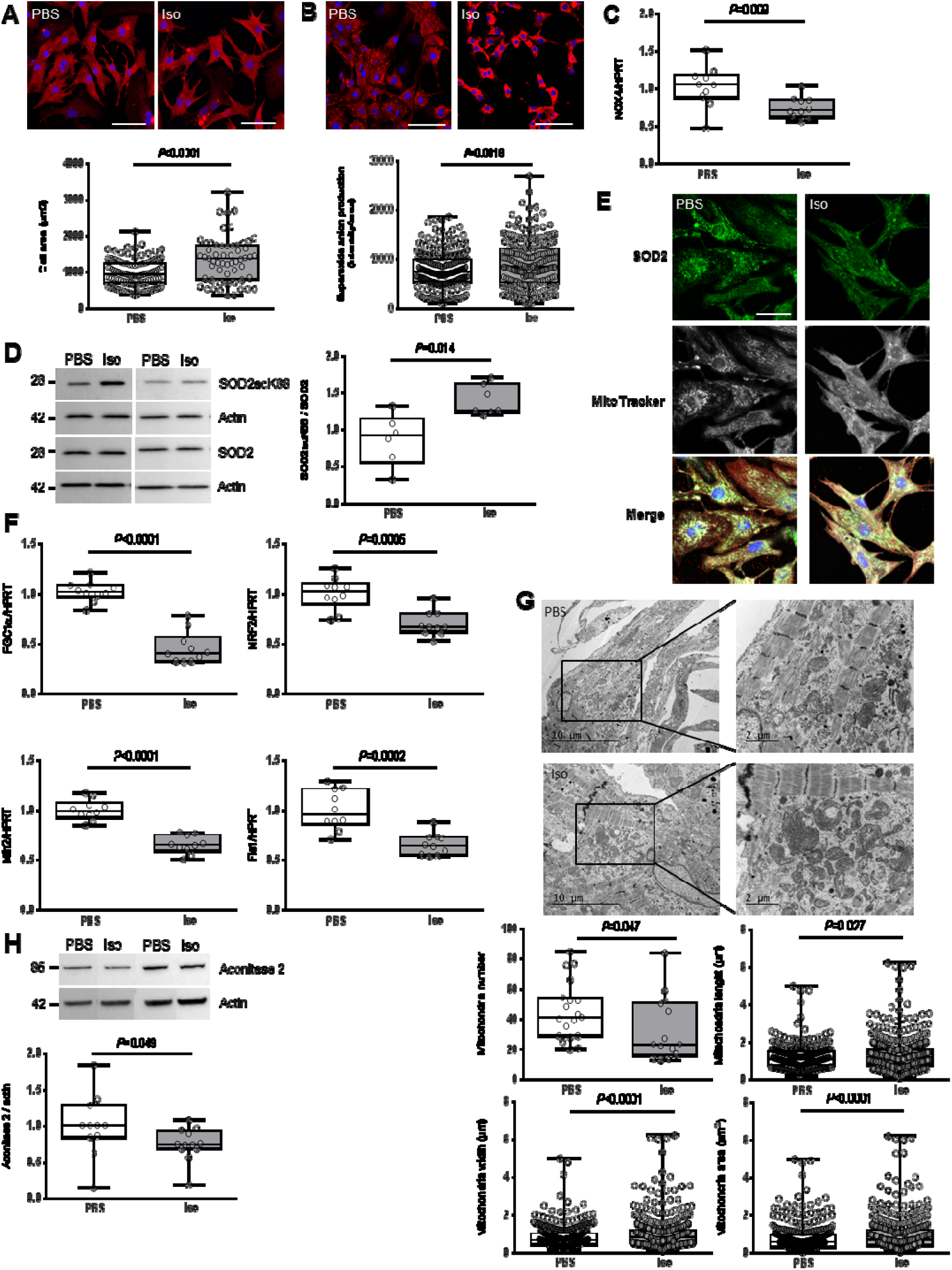
Characterization of mitochondrial oxidative stress in hypertrophied neonatal rat cardiomyocytes. (A) Hypertrophy was quantified in NCMs untreated (PBS) (white box) or treated with Isoproterenol (Iso) (grey box) for 24h by immunofluorescence of alpha-actinin (red) and nuclei (blue) (top panels) and quantification of cell area (μm^2^) (bottom panel) (from 3 independent experiments and at least 273 cells). Oxidative stress was quantified in NCMs untreated (PBS) or Iso treated for 24h by fluorescence quantification of (B) mitochondrial superoxide anion with mitoSOX probes (red) (from 3 independent experiments and at least 236 cells), by RT-qPCR of (C) NADPH oxidase (NOX) 4, and by western blot of (D) SOD2 acetylated on lysine 68 (SOD2acK68) on SOD2 ratio. (E) Representative images of SOD2 (green) localized in mitochondria (white) of NCMs untreated (PBS) or Iso treated for 24h. Colocalization appeared in merge images. Nuclei were stained by Dapi (blue). Mitochondrial biogenesis was quantified in NCMs untreated (PBS) or Iso treated for 24h by RT-qPCR of (F) peroxisome proliferator-activated receptor gamma coactivator 1-alpha (PGC1α), Nuclear Respiratory Factor (NRF) 2, mitofusin (Mfn) 2 and Fis1. Data were normalized to HPRT for RNA and actin for protein. (G) Ultrastructure of PBS and Iso treated NCMs (magnification x7000 with a scale bar of 5 μm (left images) and x12000 with a scale bar of 2 μm (right images)) and quantification of mitochondria number, length (μm), width (μm) and area (μm^2^) in NCMs untreated (PBS) or Iso treated for 24h. (H) Quantification of Krebs cycle by western blot of aconitase 2 in NCMs untreated (PBS) or Iso treated for 24h. *P* values are indicated with at least 3 independent experiments. Representative images were selected to represent the mean values of each condition.

We also quantified anti-oxidant enzymes and observed no significant modulation of catalase, superoxide dismutase 1 (SOD1) and peroxiredoxin-1 (Prx-1) (Figure S1C) due to hypertrophy. We previously showed that superoxide dismutase 2 (SOD2) was significantly increased during HF (Emilie Dubois-Deruy et al. 2017). As the lysine 68 (K68) is the most important acetylation site contributing to SOD2 inactivation (Lu et al. 2015), we quantified the acetylated form of SOD2 (SOD2acK68) and observed a significant increase of the SOD2acK68/SOD2 ratio with no modulation of total SOD2 expression (Figure 1D), meaning that SOD2 activity is significantly decreased in hypertrophied NCMs. We confirmed a mitochondrial localization of SOD2 shown by a colocalization with mitotracker probe (Figure 1E), with no impact of Iso treatment on the mitochondrial localization of SOD2 (Pearson coefficient: 0.805 for PBS, 0.806 for Iso) (Figures 1E and S1D). We also confirmed that SOD2 and SOD2acK68 are both localized in mitochondria independently of Iso treatment by subcellular fractionation (Figure S1E).

As SOD2 is a mitochondrial enzyme, we then investigated the main regulator of mitochondrial biogenesis, the peroxisome proliferator-activated receptor gamma coactivator 1-alpha (PGC1α) and its transcriptional coactivators the nuclear respiratory factor (NRF1 and NRF2) as well as different molecules involved in mitochondrial fusion and fission in hypertrophied NCMs. We first observed a significant decrease of mitochondrial biogenesis quantified by PGC1α, NRF1 and NRF2 RNA levels (Figures 1F and S1F). Moreover, RNA levels of mitofusin 2 (Mfn2) as well as mitochondrial fission 1 protein (Fis 1) were significantly decreased in hypertrophied NCMs (Figure 1F).

All these results suggested that mitochondrial biogenesis is impaired during hypertrophy, which is corroborated by electronic microscopy showing a significant decrease of the number of altered (loss of electron-dense matrix) and larger (significant increased length, width and area) mitochondria (Figure 1G). We also observed a significant decrease of aconitase 2, an enzyme of Krebs cycle reflecting mitochondrial function (Figure 1H). We then quantified mitochondrial respiration by oxygraphy using isolated NCMs treated or not with Iso. We observed a classical profile, including: basal respiration, ATP production-coupled respiration (oligomycin A), maximal and reserve capacities (carbonyl cyanide m-chlorophenyl hydrazine (CCCP)) and non-mitochondrial respiration (antimycin A (AA)) but we did not observe any significant modulation of oxygen consumption upon Iso treatment compared to untreated NCMs (Figure S1G). These results suggested that mitochondria biogenesis is strongly decreased, with no detectable impact on oxygen consumption, during cardiac hypertrophy induced by β-adrenergic stimulation.

### Characterization of mito(auto)phagy in hypertrophied NCMs

As mitophagy is the selective pathway of degradation of defective mitochondria, we investigated how hypertrophy could affect this process. As the inner mitochondrial membrane depolarization is the precursor step for mitophagy, we first used the JC-1 dye, with a green fluorescence emission for the monomeric form of the probe and a red fluorescent emission for a concentration-dependent formation of J-aggregates. We observed a significant mitochondrial depolarization, indicated by the decreased red/green ratio, in hypertrophied NCMs after 24h of Iso (Figure 2A). We then quantified the proteins involved in mitophagy/autophagy process. As previously shown (Turkieh et al. 2019), Iso treatment induced a decrease of mitophagy/autophagy with a significant decrease of parkin and LC3II/LC3I ratio (Figure 2B). No modulation of ubiquitinylated proteins and beclin-1 was observed (Figure 2B). To determine if autophagy is active, NCM were treated by bafilomycin (BAF) to inhibit the autophagosome–lysosome fusion. This inhibition induced an increase of LC3I, LC3II and LC3II/LC3I ratio, but not of parkin, showing that autophagy is decreased but still active in hypertrophied NCMs (Figure 2C).

**Figure 2:**
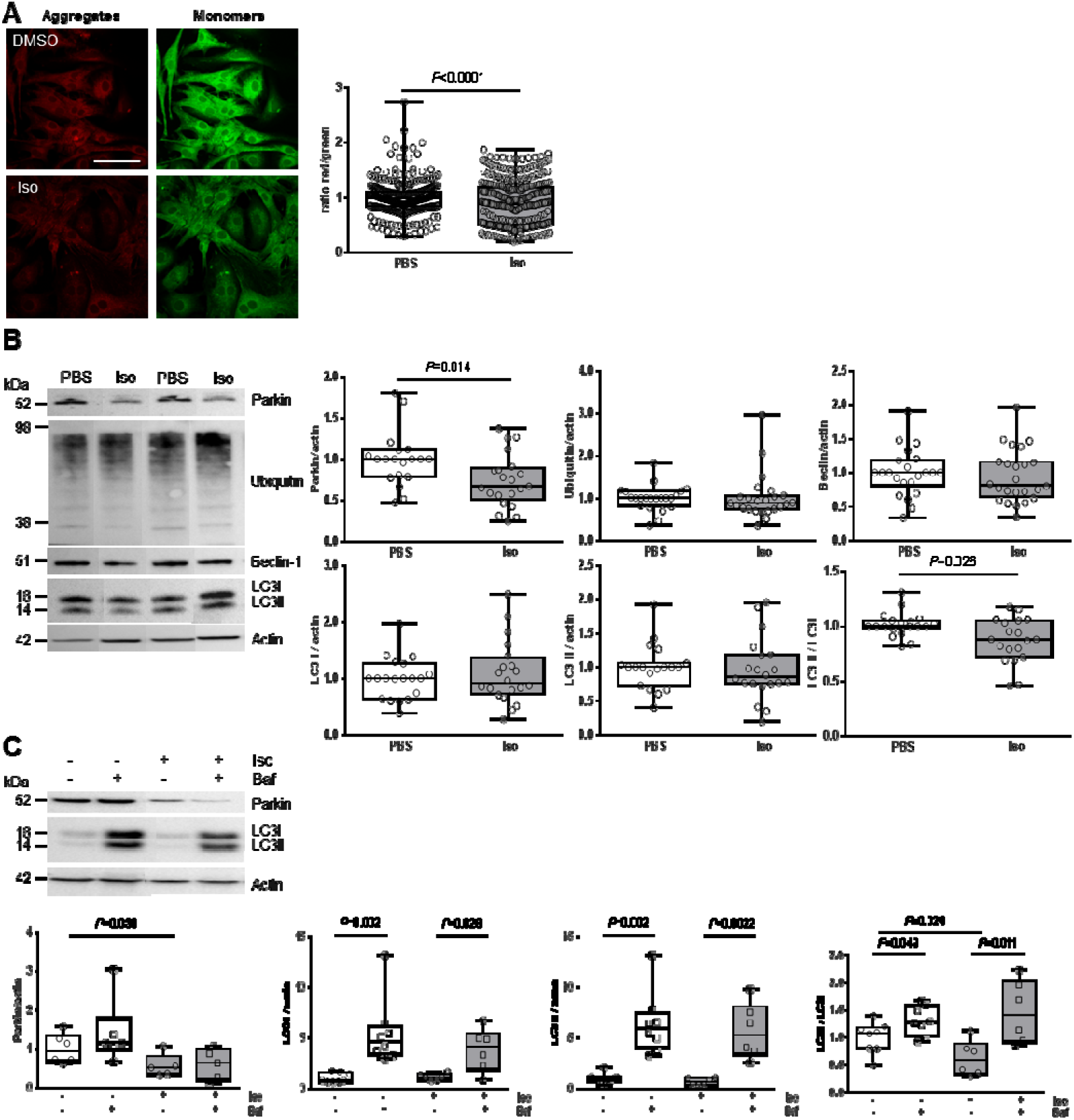
Characterization of autophagy / mitophagy in hypertrophied neonatal rat cardiomyocytes. (A) Mitochondrial membrane potential was quantified in NCMs treated with Iso for 24h by fluorescence quantification of JC-1 dye (aggregates (red) and monomer (green)) (from 3 independent experiments and at least 366 cells). Mitophagy was quantified in NCMs treated with Iso for 24h (B) and with Bafilomycin A1 (Baf) and Iso for 24h (C) by western blot of parkin, ubiquitinylated proteins, beclin-1, LC3I, LC3II and LC3II/LC3I ratio. Data were normalized to actin. *P* values are indicated with at least 3 independent experiments. Representative images were selected to represent the mean values of each condition.

### Effect of mitochondrial anti-oxidant (MitoQ) and SOD mimic (EUK 134) on oxidative stress in hypertrophied neonatal rat cardiomyocytes

To prevent cardiac hypertrophy induced by oxidative stress, we pre-treated NCMs with two antioxidants, MitoQ, a derivative of coenzyme Q that target ROS in mitochondria and the EUK 134, a mimic of SOD, non-specific of mitochondria. First, we observed two phases in cell index of NCMs pre-treated by MitoQ with an increase during the first 2h, corresponding to the pre-treatment followed by a progressive but significant decrease of cell index during the next 24h (Figure 3A). As expected, we quantified a significant decrease in cardiomyocytes area as well as in superoxide anion production in NCM pre-treated by MitoQ before Iso treatment (Figures 3B and 3C). More surprisingly, we observed a significant increase of NOX4 RNA levels in NCM pre-treated by MitoQ (Figure 3D). Interestingly, the SOD2acK68/SOD2 ratio is significantly decreased, reflecting an increase of SOD2 activity in NCMs pre-treated by MitoQ (Figure 3E). Finally, the pre-treatment did not affect SOD2 mitochondrial localization, as confirmed by colocalisation with the mitotracker probe but we observed that the mitochondria seem altered in NCMs pre-treated by MitoQ (Figure 3F). We also quantified the other anti-oxidant enzymes and observed no modulation of catalase but a significant decrease of Prx-1 and SOD1 in NCMs pre-treated by MitoQ (Figure S2A).

**Figure 3:**
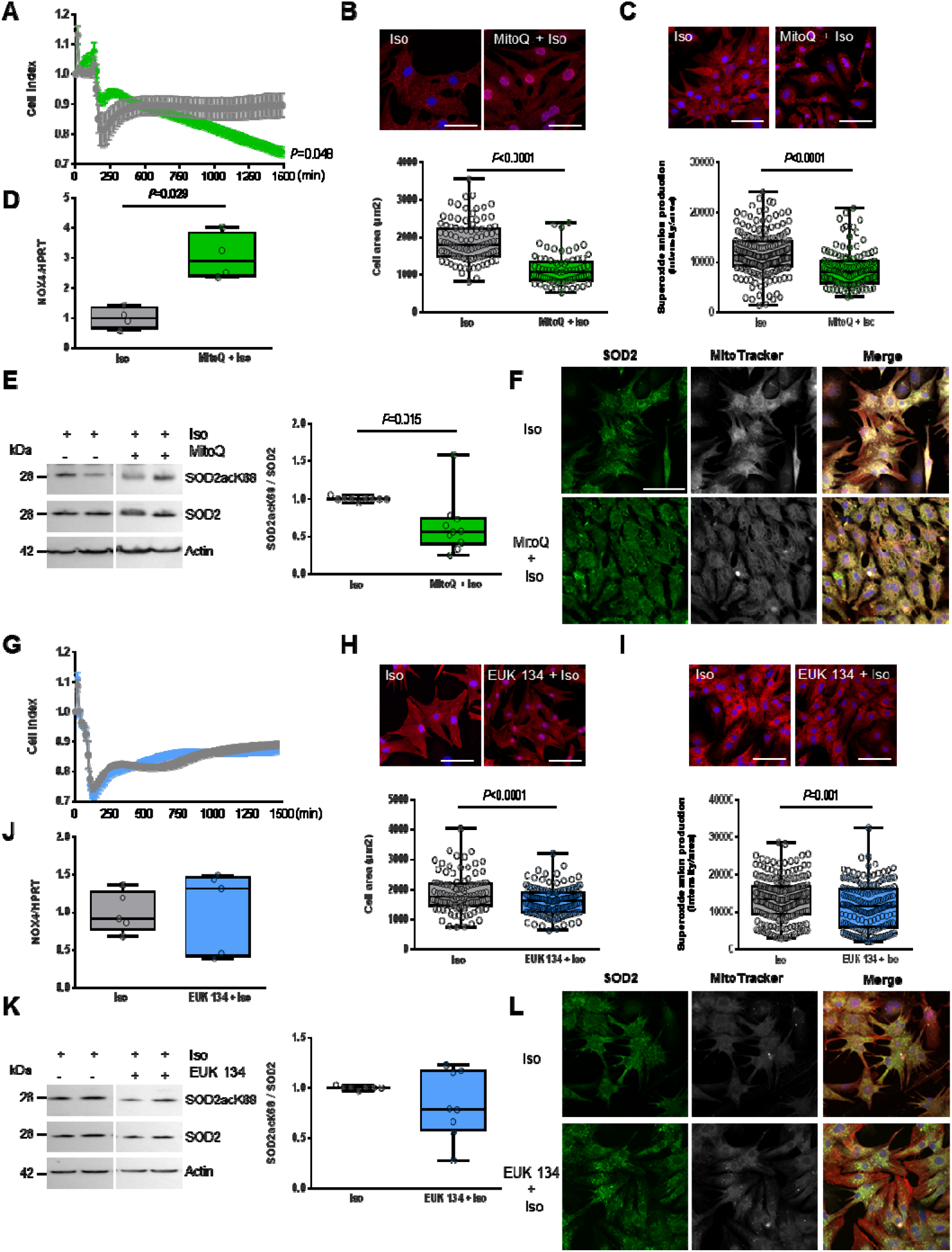
Effect of anti-oxydants in hypertrophied neonatal rat cardiomyocytes. (A and G) Cell index quantification by RTCA analysis in NCMs Iso treated (grey line) with mitoquinone (MitoQ) (green line, A) or EUK 134 pre-treatment (blue line, G). Cell index was recorded every 15 minutes (n=4 independent isolations, in duplicate) and statistical analyses were performed by functional ANOVA (R package fdANOVA). Hypertrophy was quantified in NCMs treated with Iso for 24h with or without MitoQ (B) or EUK 134 (H) pre-treatment by immunofluorescence of alpha-actinin (red) and nuclei (blue) and quantification of cell area (μm^2^) (from 3 independent experiments and at least 75 cells). Mitochondrial superoxide anion was quantified in NCMs treated with Iso for 24h with or without MitoQ (C) or EUK 134 (I) pre-treatment by fluorescence quantification of mitoSOX (red) (from 3 independent experiments and at least 133 cells), by (D and J) RT-qPCR of NOX4 and by (E and K) western blot of SOD2acK68 on SOD2 ratio. (F and L) Representative images of SOD2 (green) localized in mitochondria (white) in NCMs treated with Iso for 24h with or without MitoQ (F) or EUK 134 (L). Colocalization appeared in merge images. Nuclei were stained by Dapi (blue). *P* values are indicated with at least 3 independent experiments. Representative images were selected to represent the mean values of each condition.

On the contrary, we did not observe significant changes of cell index in NCMs pre-treated by EUK 134 compared to NCMs only treated with Iso (Figure 3G). As expected, we quantified a significant decrease in cardiomyocytes area as well as in superoxide anion production in NCM pre-treated by EUK 134 before Iso treatment (Figures 3H and 3I) without modulation of NOX4 RNA levels in NCM pre-treated by EUK 134 (Figure 3J). Contrary to MitoQ pre-treatment, the SOD2acK68/SOD2 ratio is not modulated in NCMs pre-treated by EUK 134 (Figure 3K). EUK 134 pre-treatment did not affect SOD2 mitochondrial localization, as confirmed by colocalisation with the mitotracker probe (Figure 3L). No modulation of the anti-oxidant enzymes was observed in NCMs pre-treated by EUK 134 (Figure S2B).

### Differential effect of mitochondrial anti-oxidant (MitoQ) and SOD mimic (EUK 134) on mitochondrial biogenesis in hypertrophied rat cardiomyocytes

In order to understand how the two antioxidants MitoQ and EUK 134 act differently in cardiomyocytes, we investigated the mitochondrial biogenesis and respiration in hypertrophied NCM pre-treated with MitoQ and EUK 134. Interestingly, we quantified a significant increase of both NRF1 and NRF2 without modulation of PGC1α RNA levels (Figure S2C) in NCMs pre-treated by MitoQ. Moreover, RNA levels of Mfn2 and Fis 1 were not modulated in hypertrophied NCMs pre-treated with MitoQ (Figure S2C). In the other hand, we did not observe any significant modulation of PGC1α, NRF1, NRF2, Mfn 2 and Fis 1 in NCMs pre-treated with EUK 134 (Figure S2D). All these results indicated that EUK 134 did not impact mitochondrial biogenesis conversely to the effect of MitoQ.

Finally, we quantified mitochondrial respiration and observed a significant decrease of oxygen consumption with a loss of respiration capacity (shown by no response to CCCP) in hypertrophied NCMs pre-treated with MitoQ whereas no difference of oxygen consumption was observed in NCM pre-treated by EUK 134 (Figure 4A). Moreover, electronic microscopy showed an aggravation of mitochondria alteration after pre-treatment with MitoQ as indicated by the increased mitochondria area, length and width associated with an important loss of electron-dense matrix whereas mitochondria were not affected by EUK 134 pre-treatment compared to only Iso treatment (Figure 4B). We also observed a significant and stronger decrease of aconitase 2 in hypertrophied NCMs pre-treated with MitoQ whereas the pre-treatment with EUK 134 did not modulate aconitase 2 expression (Figure 4C).

**Figure 4:**
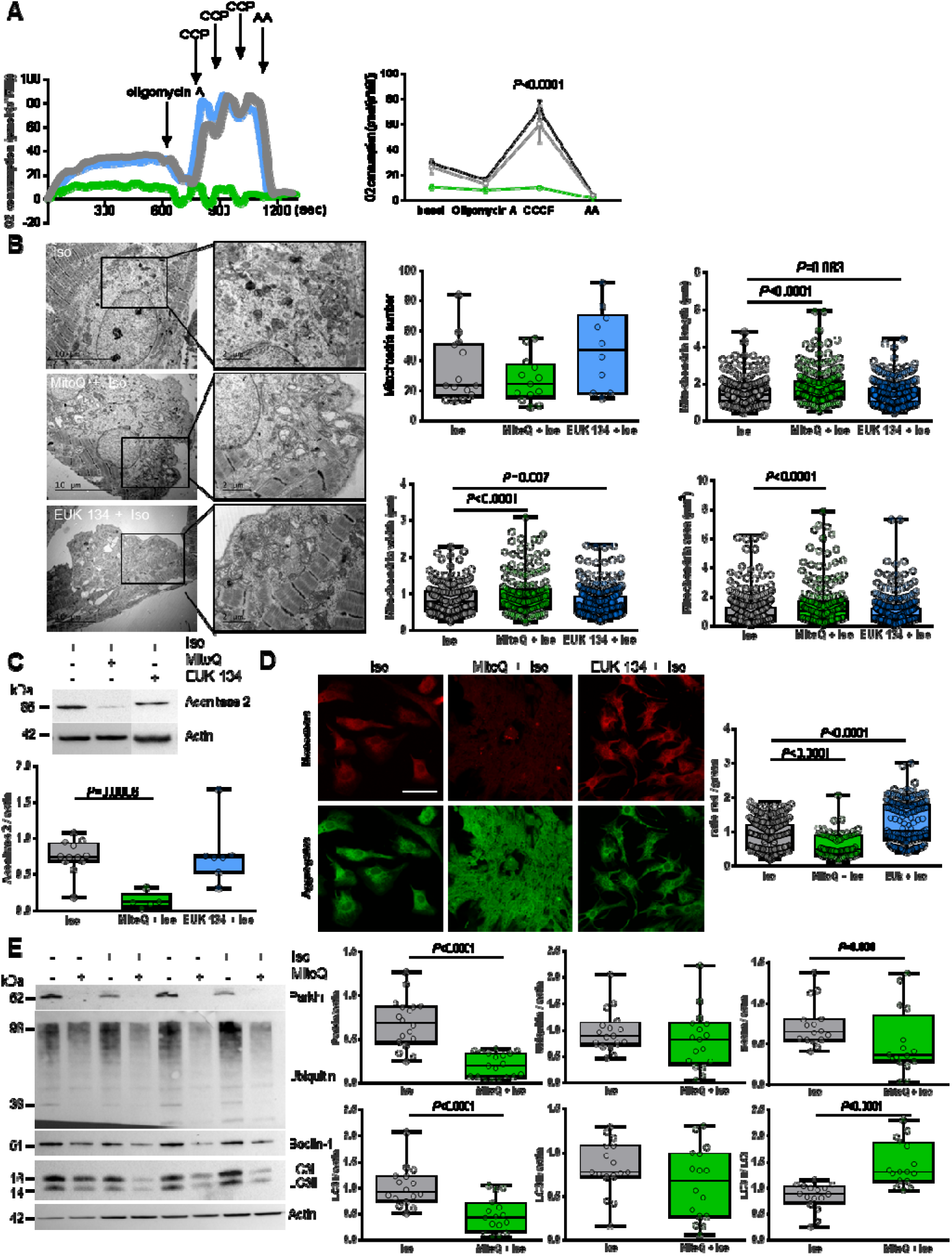
Effect of anti-oxydants on mitochondrial function and mitophagy in hypertrophied neonatal rat cardiomyocytes. Mitochondrial function was analyzed in NCMs treated with Iso for 24h with or without MitoQ or EUK-134 pre-treatment by (A) quantification of oxygen consumption, by (B) ultrastructure of NCMs (magnification x7000 with a scale bar 5 μm (left images) and x12000 with a scale bar 2 μm (right images)) and quantification of mitochondria number, length (μm), width (μm) and area (μm^2^). (C) Quantification of Krebs cycle by western blot of aconitase 2. (D) Mitochondrial membrane potential was quantified in NCMs treated with Iso for 24h with or without MitoQ or EUK-134 pre-treatment by fluorescence quantification of JC-1 dye (aggregates (red) and monomer (green)) (from 3 independent experiments and at least 94 cells). (E) Mitophagy was quantified in NCMs treated with Iso for 24h with or without MitoQ pre-treatment by western blot of parkin, ubiquitinylated proteins, beclin-1, LC3I, LC3II and LC3II/LC3I ratio. Data were normalized to actin. *P* values are indicated with at least 3 independent experiments. Representative images were selected to represent the mean values of each condition.

Of note, all these data were confirmed in NCM only pre-treated with MitoQ or EUK 134 (Figures S3A-D). These results showed that mitochondria biogenesis and function are highly altered by MitoQ, independently of Iso treatment whereas mitochondrial biogenesis and function are not affected by the anti-oxidant and anti-hypertrophic effect of EUK 134.

### Differential effect of mitochondrial anti-oxidant (MitoQ) and SOD mimic (EUK 134) on mitophagy in hypertrophied rat cardiomyocytes

As mitophagy is the selective pathway of degradation of defective mitochondria, we investigated how the two antioxidants MitoQ, and EUK 134 could affect this process. We first used the JC-1 dye and quantified a significant and stronger decrease of the red/green ratio in hypertrophied NCMs pre-treated with MitoQ whereas the pre-treatment with EUK 134 increased significantly the mitochondrial membrane potential (Figure 4D). We then quantified the proteins involved in mitophagy/autophagy process. Interestingly, parkin expression was completely suppressed by MitoQ (alone or with Iso) (Figures 4E and S3E) and MitoQ pre-treatment significantly decreased the expression of beclin-1 and LC3I without modulation of ubiquitinylated proteins (Figure 4E), suggesting a dysregulation of mitophagy/autophagy induced by MitoQ. To determine if autophagy is active, NCM were treated by bafilomycin (BAF) at the same time of MitoQ and Iso treatment. The inhibition of the autophagosome– lysosome fusion did not modulate the LC3II/LC3I ratio, suggesting that autophagy is inactive in hypertrophied NCMs pre-treated with MitoQ (Figure S3F). Conversely, EUK 134 pre-treatment did not impact mitophagy/autophagy process (Figure S3G).

### Cardiomyocyte-specificity of the impact of mitochondrial anti-oxydant (MitoQ) on mitophagy

We then, tested these two antioxidants MitoQ and EUK 134 on hypertrophied adult cardiomyocytes (ACMs). As expected, Iso induced a significant increase of cell area (Figure 5A) and mitochondrial superoxide anion (Figure 5B) in ACMs. We did not observe any modulation in cardiomyocytes area in ACMs pre-treated by EUK 134 compared to Iso alone (Figure 5A) despite a significant decrease in superoxide anion production (Figure 5B). We also quantified a significant and progressive decrease in superoxide anion production in ACMs pre-treated by MitoQ before Iso (Figure 5C). Moreover, we observed in parallel a constant decrease in cell number, probably due to strong cell mortality (Figure 5C).

**Figure 5:**
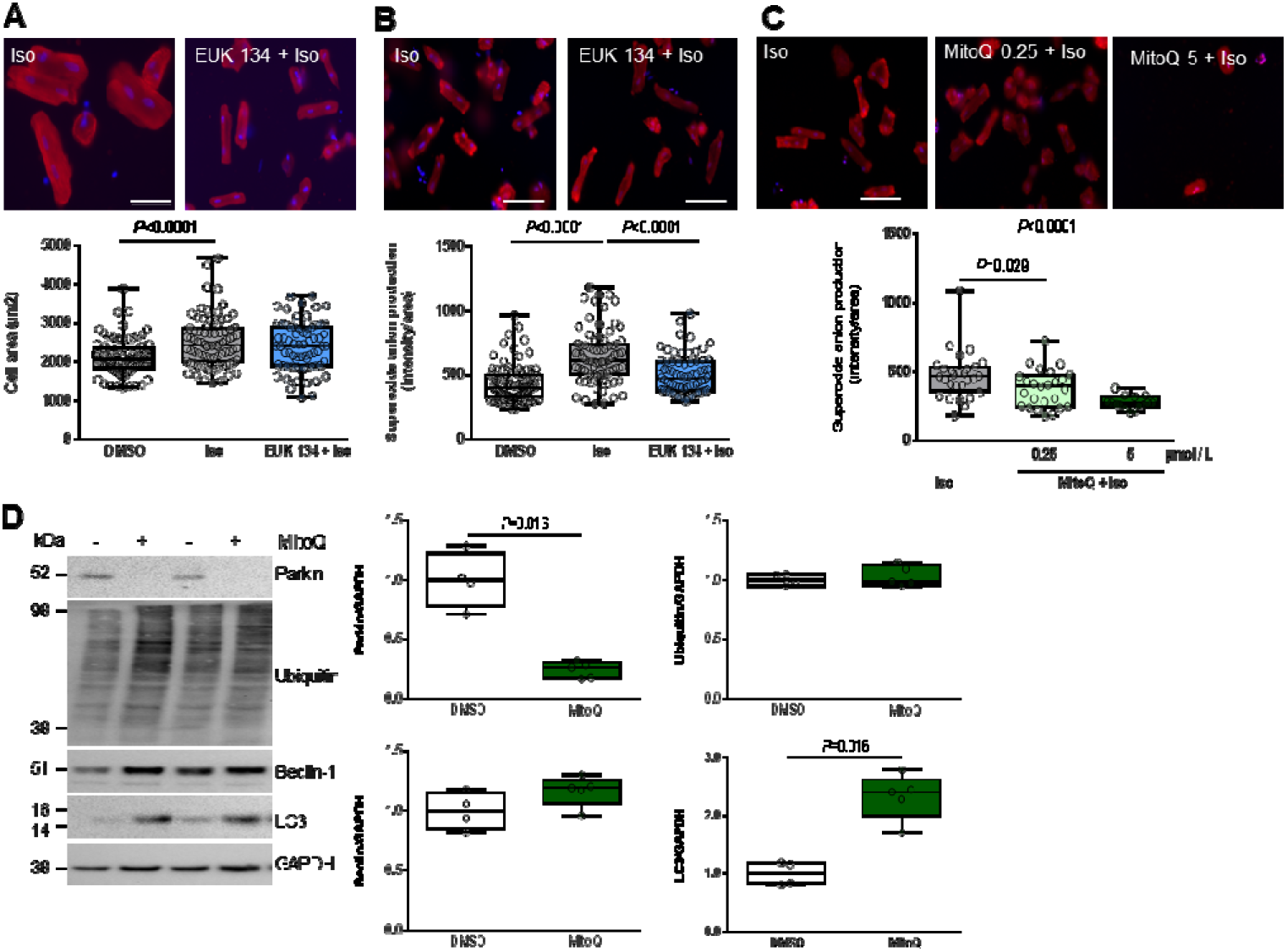
Cardiomyocytes-specificity of both anti-oxydants. (A) Hypertrophy was quantified in adult cardiomyocutes (ACMs) treated with Iso (50 μmol/L) for 48h with or without EUK 134 pre-treatment by immunofluorescence of alpha-actinin (red) and nuclei (blue) (top panels) and quantification of cell area (μm^2^) (bottom panel) (from 3 independent experiments and at least 71 cells). (B) Mitochondrial superoxide anion was quantified in ACMs treated with Iso for 48h with or without EUK 134 pre-treatment by fluorescence quantification of mitoSOX (red) (from 3 independent experiments and at least 72 cells). (C) Mitochondrial superoxide anion was quantified in ACMs treated with Iso for 48h with or without mitoQ pre-treatment (at several concentration) by fluorescence quantification of mitoSOX (red) (from at least 12 cells). (D) Mitophagy was quantified in human cardiomyocytes (HCMs) with or without MitoQ pre-treatment by western blot of parkin, ubiquitinylated proteins, beclin-1 and LC3I. Data were normalized to GAPDH. *P* values are indicated with at least 3 independent experiments. Representative images were selected to represent the mean values of each condition.

We then quantified the mitophagy/autophagy proteins in human cardiomyocytes (HCMs) and we observed a significant decrease of parkin in HCMs pre-treated by MitoQ (Figure 5D) without modulation of some proteins involved in autophagy (ubiquitinylated proteins and beclin-1). But, we observed a single band for LC3, probably LC3I, which is significantly increased in HCMs pre-treated by MitoQ as observed in NCMs (Figure 4D). In the other hand, we did not quantify any modulation of mitophagy/autophagy proteins in neonatal rat cardiac fibroblasts (NCFs) (Figure S4A), suggesting that the impact of MitoQ on mitophagy is specific of cardiomyocytes. Of note, the pre-treatment of NCFs with EUK-134 did not alter the mitophagy/autophagy process (Figure S4B).

## Discussion

In this paper, we characterized how mitochondrial oxidative stress is involved in cardiac hypertrophy and how mitochondria-targeted therapies act to prevent these effects. We also highlighted the key role of mitophagy in the deleterious effect of mitochondrial anti-oxidant (MitoQ) in mitochondrial biogenesis despite its beneficial effect by reducing mitochondrial oxidative stress and cardiomyocytes hypertrophy.

Oxidative stress is considered as a major regulator of the signal transduction in cardiac cells under pathological conditions. The understanding of the pathophysiological mechanisms which are involved in cardiac hypertrophy and remodeling process is crucial for the development of new therapeutic strategies (Rababa’h et al. 2018). Here, we used an *in vitro* model of hypertrophied cardiomyocytes (Turkieh et al. 2018) to decipher the impact of oxidative stress. We observed an increase in mitochondrial oxidative stress associated with mitochondrial dysfunction. Indeed, excessive ROS production with mitochondrial dysfunction have been described to induce irreversible damage to mitochondria, leading to the development of cardiovascular diseases (Bhatti, Bhatti, and Reddy 2017). For example, an increase in mitochondrial ROS production has been described in a murine model of MI induced by 4 weeks of coronary ligation (Ide et al. 2001) and ROS-generated by angiotensin II stimulation induced mitochondrial dysfunction, cardiomyocytes hypertrophy and HF (Dai et al. 2011). Moreover, mitophagy, the selective autophagic removal of mitochondria, is essential for clearing away the defective mitochondria but can also lead to cell damage and death if excessive. At cardiovascular levels, mitophagy is involved in metabolic activity, cell differentiation, apoptosis and other physiological processes as reviewed recently (Morales et al. 2020).

Here, we showed that mitochondrial oxidative stress is associated with an inactivation of SOD2 by acetylation. SODs are metalloproteins able to catalyze the transformation of superoxide anion into hydrogen peroxide and it is the most effective antioxidant enzyme in humans (Emilie Dubois-Deruy et al. 2020). The acetylation of K68 site is the most important acetylation site contributing to SOD2 inactivation (Lu et al. 2015) and contributes to induced hypertension (Dikalova et al. 2017). Inhibition of SOD2 expression induced both mitochondrial oxidative stress and hypertrophy of cardiomyoblasts (Emilie Dubois-Deruy et al. 2017) and mice deficient in SOD2 die of cardiomyopathy within 10 days of birth whereas the heterozygous SOD2 (+/−) mice show ultrastructural damage of the myocardium associated with an increased oxidative stress (Strassburger et al. 2005).

It was suggested that mitochondria-targeted therapies could be effective in HF (Sabbah 2016; Senoner and Dichtl 2019). In this context, we selected two anti-oxidants molecules, one targeting the mitochondria, the mitochondrial derivative of coenzyme Q (MitoQ) and one anti-oxidant with superoxide dismutase activity (EUK 134), whose pre-treatment decreased Iso-induced hypertrophy and mitochondrial oxidative stress. Surprisingly, we observed a deleterious effect of MitoQ on mitochondrial function and mitophagy. Controversial data described a protective or deleterious role of MitoQ. Indeed, *in vitro*, MitoQ has been described to prevent oxidative stress and alterations in mitochondrial proteins observed in palmitic acid -stimulated cardiomyoblasts (Jiménez-González et al. 2020). *In vivo*, MitoQ reduced cardiac oxidative stress and prevented the development of cardiac fibrosis and hypertrophy in obese rats (Jiménez-González et al. 2020). MitoQ could also significantly improved LV dysfunction and increased metabolism-related gene expression in mice subjected to ascending aortic constriction (Kim et al. 2020). These discrepancies could be explained by the cell-specificity. Indeed, here, we observed specifically a decrease in parkin expression in human and rat cardiomyocytes but not in the fibroblasts. Nevertheless, according to our data, some studies, notably in cancer cells, described that MitoQ could lead to ROS production, rapid membrane depolarization and apoptotic cell death (Doughan and Dikalov 2007; Pokrzywinski et al. 2016). Indeed, MitoQ could cause mitochondrial swelling and depolarization in kidney proximal tubule cells by a mechanism non-related to the anti-oxidant activity but most likely because of the increased inner mitochondrial membrane permeability due to insertion of the alkyl chain (Gottwald et al. 2018). These side effects of MitoQ are in accordance with our data with impaired cardiomyocytes respiration, in link with the mitochondria alteration shown by their electronic microscopy ultrastructure and membrane potential. The more striking point is the abolished expression of parkin upon MitoQ treatment leading to defective mitophagy and accumulation of deficient mitochondria in cardiomyocytes.

Interestingly, the other anti-oxidant tested, EUK 134 seems more promising with no obvious side effects. Until now, EUK 134 was described to decrease hypertrophy and oxidative stress and restore mitochondrial membrane potential in cardiomyoblasts H9c2 hypertrophied by phenylephrine (Purushothaman and Nair 2016). *In vivo*, EUK 134 improved remodelling induced by hindlimb unloading as well as atrophy of muscle fibers by decreasing ROS production (Kuczmarski et al. 2018) and attenuated cardiomyocyte hypertrophy, oxidative stress, proapoptotic signalling and interstitial fibrosis in congestive HF in the rat monocrotaline model of pulmonary arterial hypertension. Here, this SOD mimic restored impaired cardiomyocyte metabolism and mitochondrial biogenesis without affecting mitophagy. Its effect on mitochondrial oxidative stress was observed both in neonatal and adult cardiomyocytes.

In conclusion, anti-oxidant therapeutic strategies should take into account the functional interaction between mitochondrial dynamism, biogenesis and mitophagy (Song et al. 2017).

The new hypothesis related to this work are that both anti-oxidant (MitoQ and EUK 134) improve cardiac hypertrophy and ROS production in cardiomyocytes but MitoQ induces impaired mitochondria morphology, oxygen respiration and membrane potential through a defect in mitophagy, specifically in cardiomyocytes whereas EUK 134 restores impaired cardiomyocyte metabolism and mitochondrial biogenesis.

## Methods

### Cell Culture

#### Primary cultures of neonatal rat cardiomyocytes and fibroblasts

Primary cultures of NCMs and fibroblasts (NCFs) were prepared from heart ventricles of 1-or 2-day-old rats, killed by decapitation as previously described (Bouvet et al. 2016; E. Dubois-Deruy et al. 2015). Briefly, cardiac cells were dissociated by enzymatic digestion with 0.04% collagenase II (Worthington, Lakewood, NJ, USA) and 0.05% pancreatin (Sigma-Aldrich) at 37°C. NCMs and NCFs were removed from cell suspension by centrifugation 30 min at 3000 rpm in a discontinuous Percoll gradient (bottom 58.5%, top 40.5%, Sigma-Aldrich).

NCMs were seeded at a density 4 or 8 × 10^5^ cells/well in 6-well plates coated with 0.01% of collagen (Sigma-Aldrich) and cultured in a medium containing DMEM/Medium199 (4:1), 10% horse serum (Life Technologies), 5% fetal bovine serum (ATCC), 1% penicillin and streptomycin (P/S) (10,000 U/mL, Life Technologies) at 37°C under 5% CO_2_ atmosphere. NCMs were starved for 24h before Isoproterenol (Iso, 10 μmol/L) or PBS (as control) treatment for 24h. A pre-treatment of mitoquinone (MitoQ, 1 μmol/L, 2h), EUK 134 (10 μmol/L, 1h) or DMSO (as control) were also used. For autophagy experiments, Bafilomycin A (Baf, 10 nmol/L) could also be added during 24h at the same time of the pre-treatment with both anti-oxidants and the totality of Iso treatment.

NCFs were seeded at a density 3.5 × 10^5^ cells/well in 6-well plates and cultured in a medium containing DMEM/Glutamax, 10% fetal bovine serum (ATCC), 1% P/S (10,000 U/mL, Life Technologies) at 37°C under 5% CO_2_ atmosphere. NCFs were starved for 24h before pre-treatment of mitoquinone (MitoQ, 1 μmol/L, 2h), EUK 134 (10 μmol/L, 1h) or DMSO (as control). Cells were then maintained during 24h in the privation medium.

#### Primary cultures of adult rat cardiomyocytes

Adult rat cardiomyocytes (ACMs) were isolated as previously described (Bertrand et al. 2006). Briefly, hearts were collected from male Wistar rats (250g) and perfused in Krebs-Henseleit buffer containing 10 mmol/L HEPES, 5 mmol/L glucose, 2 mmol/L pyruvate and 25 mmol/L NaCl. Hearts were then digested in Krebs-Henseleit buffer containing 1mg/mL collagenase type II and 0.4% free fatty acid BSA for 30 min at 37°C. After collagenase perfusion, hearts were removed from the perfusion apparatus and cut in small fragments. They were then incubated under agitation for 10 min at 37°C in Krebs-Henseleit Buffer containing 0.02 mmol/L CaCl_2_ and calcium chloride was progressively added to the medium to reach 1 mmol/L final concentration. After sedimentation and washing, cells were finally resuspended in MEM medium containing 20 mmol/L HEPES, 2.5% fetal bovine serum and 2% penicillin / streptomycin and equally seeded in 3-cm diameter wells (20 wells for one digested heart) previously coated with laminin and incubated at 37°C for at least 1h. Two hours after seeding, ACMs were starved for 24h before Isoproterenol (Iso, 50 μmol/L) or PBS (as control) treatment for 48h. A pre-treatment of mitoquinone (MitoQ, 0.25 to 5 μmol/L, 2h), EUK 134 (10 μmol/L, 1h) or DMSO (as control) were also used.

#### Human cell line

Human Cardiac Myocytes (HCMs) (C-12810, PromoCell) were seeded at a density 1.5 × 10^5^ cells/well in 12-well plates and cultured in a myocyte growth medium (C-22170, PromoCell) containing 0.5 ng/ml epidermal growth factor, 2 ng/ml basic fibroblast growth factor, 5 μg/ml insulin and 5% fetal calf serum. HCMs were starved for 24h before pre-treatment of mitoquinone (MitoQ, 1 μmol/L, 2h), EUK 134 (10 μmol/L, 1h) or DMSO (as control). Cells were then maintained during 24h in the privation medium.

### RNA Extraction and qRT-PCR analyses

#### RNA extraction and qRT-PCR analyses

RNA was extracted from NCMs with QIAGEN RNeasy Mini Kit (Qiagen), as described by the manufacturers’s instructions. Reverse-transcription was performed with 100, 250 or 500 ng of total RNA using the miScript II RT kit (Qiagen) and the cDNA was amplified with miScript SYBR Green PCR (Qiagen) on an Aria Mx Q-PCR system (Agilent Technologies), according to the manufacturer’s instructions. The sequences of the different primers (Eurogentec) used were: Fis1 (sense: GCACGCAGTTTGAATACGCC, antisense: CTGCTCCTCTTTGCTACCTTTGG); hypoxanthine Phosphoribosyltransferase 1 (HPRT) (sense: ATGGGAGGCCATCACATTGT, antisense: ATGTAATCCAGCAGGTCAGCAA); Mfn2 (sense: GATGTCACCACGGAGCTGGA, antisense: AGAGACGCTCACTCACTTTG); NOX4 (sense: ATCTGGGTCTGCAGAGACAT, antisense: CTGAGGTACACTGATGTT); NRF1 (sense : CGCAGTGACGTCCGCACAGA, antisense: AAGGTCCTCCCGCCCATGCT), NRF2 (sense: GCAACTCCAGAAGGAACAGG, antisense: AGGCATCTTGTTTGGGAATG) and PGC1α (sense: AAAAGCTTGACTGGCGTCAT, antisense : TCAGGAAGATCTGGGCAAAG). ∆∆CT method was used for data analysis.

### Protein extraction and western blot

#### Protein extraction

Proteins were extracted from 6-well plates cell into ice-cold RIPA buffer (50 mmol/L Tris [pH7.4], 150 mmol/L NaCl, 1% Igepal CA-630, 50 mmol/L deoxycholate, and 0.1% SDS) containing anti-proteases (CompleteTM EDTA-free, Roche Diagnostics), serine/threonine protein phosphatase inhibitors (Phosphatase inhibitor Cocktail 3, Sigma-Aldrich) and 1 mmol/L Na_3_VO_4_. Lysates were incubated for 1h at 4°C, centrifuged 15 min at 11,000 g to collect the soluble proteins. Protein concentrations were determined with a Lowry-based method protein assay (Biorad, Marnes-la-Coquette, France) and samples were kept at −80°C.

#### Cytosol-mitochondria fractionation

NCMs were seeded at 2,000,000 cells/dish before Iso treatment as described above. After treatment, media was removed and cells washed with 5 mL ice-cold PBS and scrapped in 750 μL of ice-cold buffer H (0.3 mol/L sucrose, 5 mmol/L TES, 2 mmol/L EGTA, pH 7.2) (Fazal et al. 2017). Scrapped cells were then transferred in a 2 mL tube in which 500 μL of buffer H containing 1 mg/mL BSA was added. After 6 times inversion, the tubes were centrifuged for 10 min at 500 g and the supernatant was collected before 10 min centrifugation at 3,000 g. Cytoplasmic fraction (supernatant) was collected and 100 μL of RIPA 2X buffer was added. Mitochondrial fraction (pellet) was resuspended in 50 μL RIPA 2X buffer.

#### Western blot

Soluble proteins (10 to 50 μg) were resolved on NuPAGE 4-12% Bis-Tris Protein Gels (Life Technologies) or on SDS-PAGE gels (12 or 15%, depending on the proteins analysed) and transferred on 0.2 μm nitrocellulose membranes (Trans-Blot^®^ Turbo^TM^ Transfert Pack, Bio-Rad). Equal total proteins loads were confirmed by Ponceau red (0.1% Ponceau, Sigma-Aldrich), 5% acetic acid (v/v)] staining of the membranes. The membranes were then blocked in 5% milk or 5% BSA in TBS-Tween buffer for 1h before 4°C overnight incubation with primary antibodies diluted in blocking solution. Blots were then washed three times with TBS-Tween 0.1 % buffer and incubated with corresponding secondary antibodies for 1h (1/5,000 to 1/10,000) in blocking solution. The Chemidoc^®^ camera (Biorad) was used for imaging and densitometry analysis after membranes were incubated with enhanced chemiluminescence (ECL™) western blotting detection reagents (GE Healthcare).

#### Antibodies

The primary antibodies used for western blot analysis were: aconitase 2 (Gentex, GTX109736), ATP synthase alpha (A-21350, Invitrogen, 1/5000), beclin-1 (3738, Cell signaling, 1/1000); catalase (CO979, Sigma-Aldrich, 1/1000); GAPDH (sc-365062, Santa Cruz, 1/10000), LC3 B (2775, Cell signaling, 1/2000); parkin (702785, ThermoFisher, 1/1000); peroxiredoxin-1 (prx1) (ab59538, Abcam, 1/1000); sarcomeric-actin (M0874, Dako, 1/5000), SOD1 (10269-1-AP, proteintech, 1/1000), SOD2 (ab 13533, Abcam, 1/5000), SOD2 acetylated on lysine 68 (SOD2acK68) (ab 137037, Abcam, 1/1000), ubiquitinylated proteins (BML-PW8810-0500, Enzo Life Science, 1/2500). The horseradish peroxidase-labelled secondary antibodies used were: anti-rabbit IgG (NA934V) and anti-mouse IgG (NA931) antibodies from GE healthcare.

### Transmission electronic microscopy

NCMs were fixed overnight in 2.5% glutaraldehyde in 0.1 mol/L phosphate buffer, and then washed 3 times with Phosphate buffer 0.1 mol/L. Samples were then post-fixed in 1% osmium tetroxide in Phosphate buffer 0.1 mol/L at room temperature for 1h then followed by dehydration steps (5 min in ethanol 50%, 5 min in ethanol 70%, 5 min in ethanol 80%, 2 x 15 min in ethanol 95%, 3 x 20 min in ethanol 100%). Cells were then detached and centrifuged 10 min at 12,000 rpm. Pellets were then incubated with propylene oxide for 30 min. Samples were then stained in propylene oxide/Epon (V/V) for 1h, then in Epon 100% twice for 1h, followed by overnight incubation before capsules embedding at 60°C for 4 days.

Ultrathin sections (85 nm) were performed with a Leica UM EC7 ultramicrotome, and sections were contrasted by uranyl acetate 2%/ethanol 50% treatment for 8 min followed by Reynolds lead citrate for 8 min. Sections were observed using a Zeiss EM900 electron microscope with Gatan Orius SC1000 camera. Quantification of mitochondria number, length, width and area were performed with analysis software (Zeiss).

### Immunofluorescence

Biphotonic confocal microscopy was used for the imaging of 4% paraformaldehyde and 0.1% Triton fixed/permeabilized cardiomyocytes. Immunofluorescence staining was performed by saturation for 30 min with 1% BSA before incubation with anti-α-actinin antibody (A-7811, Sigma-Aldrich) or anti-SOD2 (ab 16956, Abcam) (dilution 1/50) overnight at 4°C. Alexa

Fluor^®^ 568 coupled anti-mouse secondary antibody at dilution 1/300 was incubated for 30 min at room temperature before nuclei staining for 10 min at room temperature (Hoechst 33258, Invitrogen™) with mounting medium (Vectashield).

Cardiomyocytes were incubated in Hank’s Balanced Salt Solution (HBSS) with 10 μmol/L MitoSOX Red (ThermoFisher Scientific) to stain mitochondrial superoxide anion levels and 100 nmol/L Mitotracker Deep red (M22426, ThermoFisher Scientific) to stain mitochondria or 10 μg/mL of JC-1 sensor to quantify the mitochondrial membrane potential for 30 min at 37°C.

Stainings were visualized with a x40 objective on an LSM710 confocal microscope that used Zen image acquisition and analysis software (Zeiss). Images were acquired with a resolution of at least 1024×1024 and analysed with Image J software.

### Detection of mitochondrial hydrogen peroxide levels using MitoPY1

NCMs were seeded at 20,000 cells/well in 96 well-plate. The cells were serum-deprived for 24h and treated with PBS (control) or Iso at 10 μmol/L for 24h in serum-free medium. After treatment, media was removed and cells were washed twice with 100 μL PBS 1X/well and then immediately stained with 50 μmol/L MitoPY1 probe (4428, Tocris) for 30 min at 37°C in PBS 1X. After staining, cells were washed twice in 100 μL PBS 1X. Then 200 μL PBS 1X was added in 96 well-plates and absorbance was measured with a microplate reader at an excitation wavelength of 485 nm and an emission wavelength of 520 nm.

### Oxygraphy analysis

NCMs were seeded at 400,000 cells/well in 6 well-plate (1 plate per condition) in culture medium with serum. Then, cells were serum-deprived for 24h and treated with PBS (control) or Iso at 10 μmol/L for 24h in serum-free medium. A pre-treatment of MitoQ (1 μmol/L, 2h), EUK 134 (10 μmol/L, 1h) or DMSO (as control) were also used. At the end of treatment, NCMs were gently rinsed twice with PBS. Pre-warmed trypsin solution was adding in each well and incubated at 37°C for 10 min. Once cells appear detached, trypsin was inactivating by addition of medium containing serum and centrifuged at 800 g for 7 min. After removing the supernatant, the cell pellet was resuspended in pre-warmed medium with serum and count with Malassez cell.

NCM (between 1 and 1.5 million) were incubated into the O2K oxygraph chambers (Oroboros Instruments, Innsbruck, Austria) at 37°C under constant stirring. After a 15 min stabilization leading to resting respiration, oligomycin A (5 nmol/L) was added to measure leak respiration for 5 min. Then, CCCP pulses (0.5 to 2.5μmol/L steps) were performed until maximal oxygen consumption was achieved. Non-mitochondrial oxygen consumption was obtained after AA (2.5 μmol/L) injection for 5 minutes.

### Cell index quantification by RTCA

NCMs were seeded in 0.64 cm² well covered with 80% gold electrodes (00300600840, ACEA Biosciences) from which a low voltage alternating current (20 mV) is generated in the culture medium from one electrode to another by an iCELLingence system (00380601000, ACEA Biosciences). The electrical resistance of adherent cells (RCell) is registered by a control unit (00380601430, ACEA Biosciences) equipped with RTCA Lite software (00310100210, ACEA Biosciences) and is expressed at each time in cellular index (ICell) defined by the equation: ICell = (RCell - Rm) / Rm where Rm corresponds to the resistance of the medium without the cells. The L8 E-Plates are incubated at 37°C and 5% CO_2_ and the cell index is recorded every 15 min until the end of the experiment.

### Statistical analysis

Data are expressed as medians with interquartile ranges and analysed with GraphPad software version 7.0. Data were compared using nonparametric Mann–Whitney test. For RTCA experiments, the data were analysed by functional ANOVA, performed using the R package fdANOVA (Górecki and Smaga 2019). Statistical significance was accepted at the level of *P*<0.05.

## Abbreviations

SOD2acK68: Acetylated form of SOD2 in lysine 68
ACMs: adult rat cardiomyocytes
AA: antimycin A
CCCP: carbonyl cyanide m-chlorophenyl hydrazine
HF: heart failure
HCMs: human cardiac myocytes
Iso: isoproterenol
HPRT: hypoxanthine phos^horibosyl transferase
Mfn2: mitofusin 2
Mfn2: mitochondrial fission 1 proteinw
MitoQ: mitoquinone
NCFs: rat neonatal cardiac fibroblasts
NCMs: neonatal rat cardiac myocytes
NOX4: NADPH Oxidase 4
NRF: nuclear respiratory factor
PGC1α: peroxisome proliferator-activated receptor gamma coactivator 1-alpha
Prx-1: peroxiredoxin-1
ROS: reactive oxygen species
RTCA: real time cell analysis
SOD: superoxide dismutase

## Acknowledgments

We thank Dr N. Malmanche for confocal microscopy and the UMS 2014 – US 41 – PLBS – Plateformes Lilloises en Biologie & Santé for cell imaging and animal facility. We would like to thank Servier for the network used in the graphical abstract (Servier Medical Art https://smartservier.com).

## Authors contributions

EDD designed the study, make experiments and wrote the paper. VP, OB, SL, LBu, NF, AL and MC make experiments. LBe contributed to experimental models. PA and FP designed the study and wrote the paper.

---

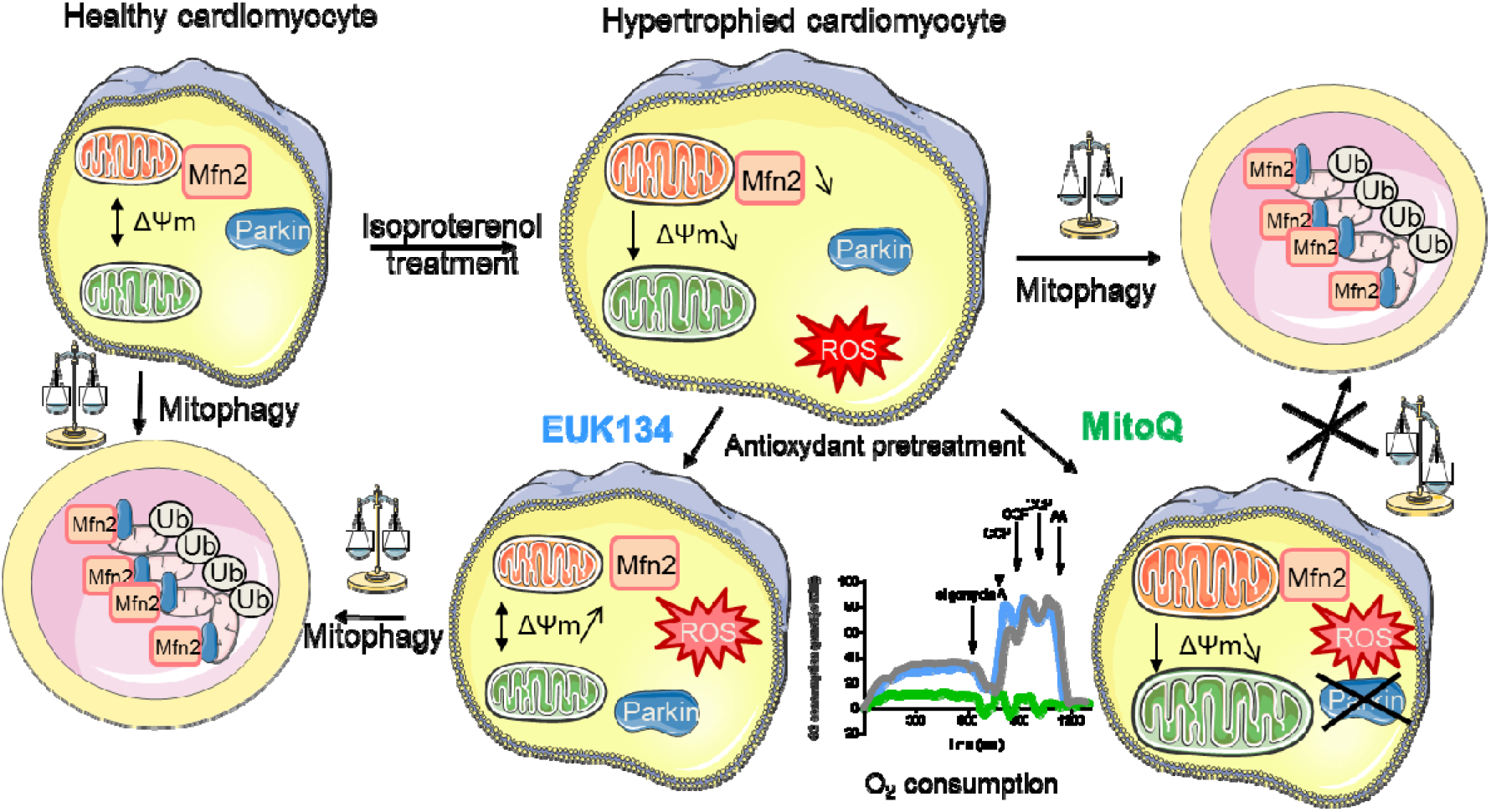
Graphical abstract.

## Supplemental Figure legends

**Figure S1:**
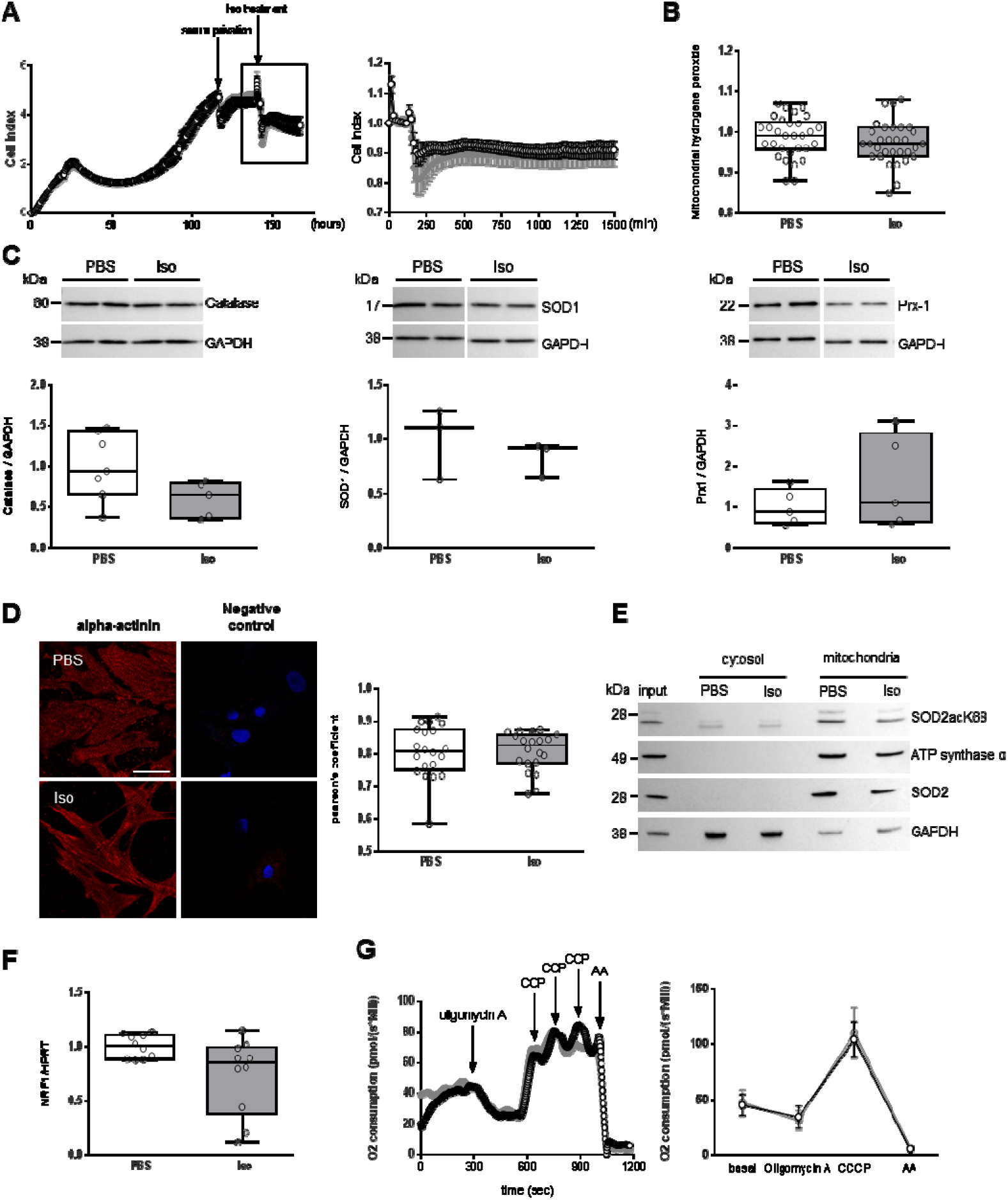
Characterization of mitochondrial oxidative stress in hypertrophied neonatal rat cardiomyocytes. (A) Cell index quantification of untreated-(black line) or Isoproterenol (Iso) treated (grey line) -NCMs by RTCA analysis. Cell index was recorded every 15 minutes for 6 days (n=4 independent isolations, in duplicate) and expressed as arbitrary unit (left panel). The square represents the curve from Iso treatment, zoomed in right panel. Statistical analyses of cell index were performed by functional ANOVA (R package fdANOVA). Oxidative stress was also quantified in NCMs untreated (PBS) or Iso treated for 24h by mitochondrial hydrogen peroxide quantification (B), and by western blot of (C) catalase, superoxide dismutase 1 (SOD1) and peroxiredoxin-1 (Prx1). (D) Colocalization was quantified by pearson’s coefficient, describing the correlation between the intensities of SOD2 and mitotracker images with the JACoP puglins on ImageJ software. (E) Representative images of SOD2acK68 and SOD2 localized in mitochondria (validated by ATP synthase α) after sub-cellular fractionation in NCMs untreated (PBS) or Iso treated for 24h. (F) Mitochondrial biogenesis was quantified in NCMs untreated (PBS) or Iso treated for 24h by RT-qPCR of Nuclear Respiratory Factor (NRF) 1. (G) Quantification of oxygen consumption at basal level and after oligomycin, carbonyl cyanide m-chlorophenyl hydrazine (CCCP) and antimycin A (AA) addition to characterize mitochondrial respiration. Data were normalized to HPRT for RNA and GAPDH or actin for protein and are presented as box-and-whisker plots showing median (line) and interquartile ranges (IQR). Representative images were selected to represent the mean values of each condition.

**Figure. S2:**
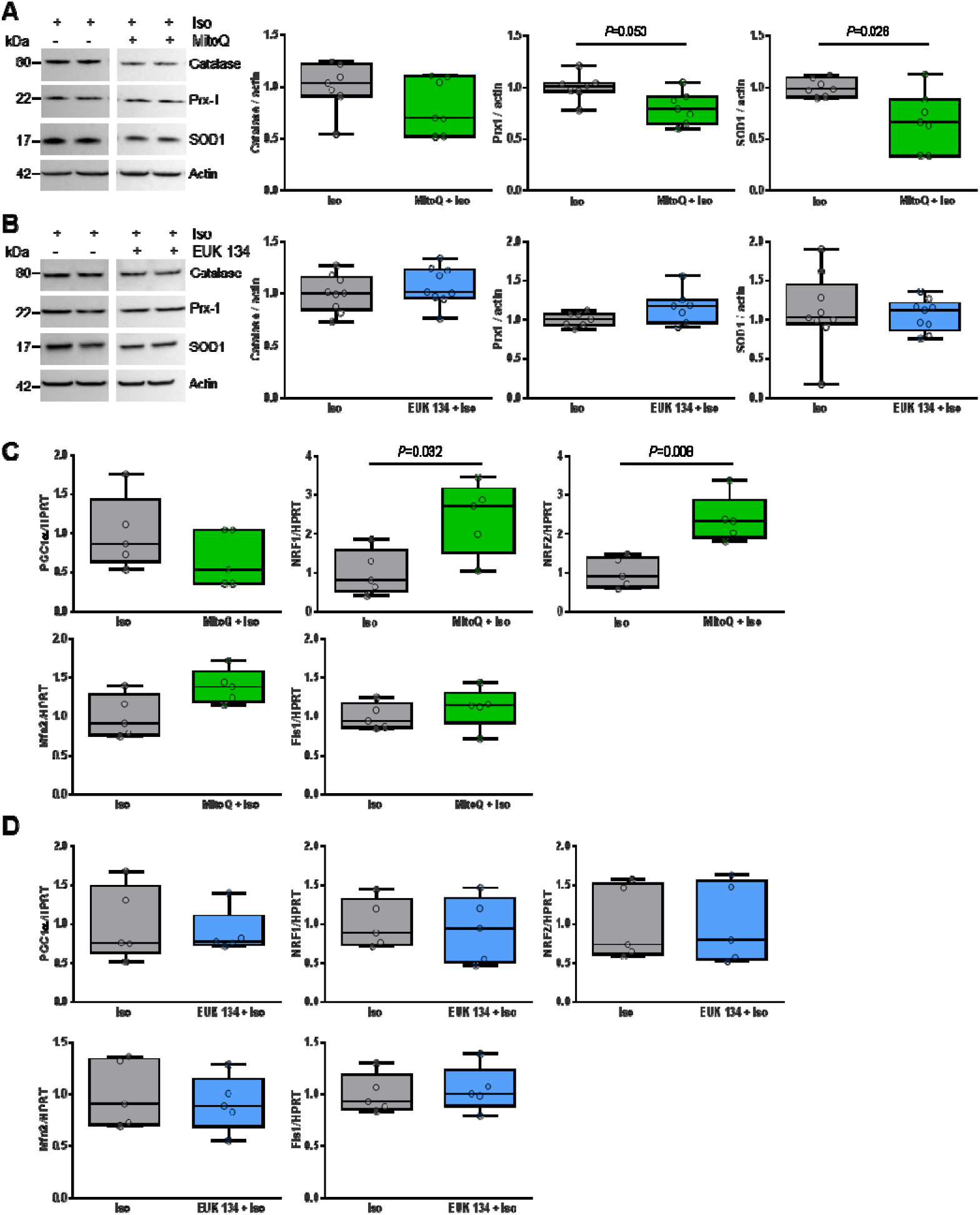
Effect of anti-oxydants (mitoquinone and EUK 134) on hypertrophied neonatal rat cardiomyocytes. Oxidative stress was quantified in NCMs Iso treated with or without MitoQ (A) or EUK 134 (B) pre-treatment by western blot of catalase, Prx1 and superoxide dismutase 1 (SOD1). Mitochondrial biogenesis was quantified in NCMs Iso treated with or without MitoQ (C) or EUK 134 (D) pre-treatment by RT-qPCR of peroxisome proliferator-activated receptor gamma coactivator 1-alpha (PGC1α), Nuclear Respiratory Factor (NRF) 1 and 2, mitofusin (Mfn) 2 and of Fis1. *P* values are indicated with at least 3 independent experiments. Representative images were selected to represent the mean values of each condition.

**Figure S3:**
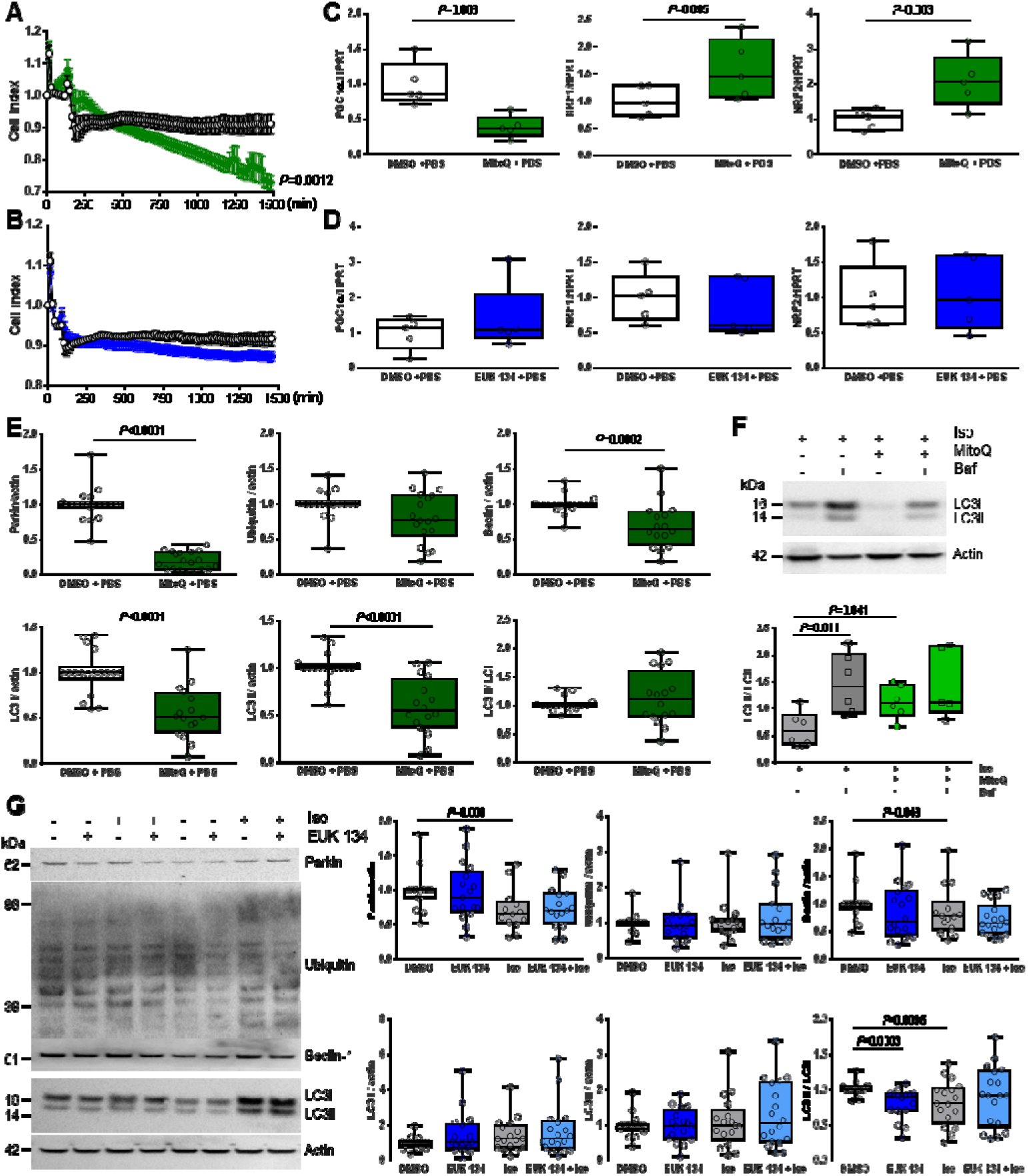
Effect of both anti-oxydants on oxidative stress, mitochondrial biogenesis and mitophagy in hypertrophied neonatal rat cardiomyocytes. (A and B) Cell index quantification by RTCA analysis in NCMs controls (DMSO + PBS) (black line) treated with mitoquinone (MitoQ) (green line) pre-treatment (A) or with EUK 134 (blue line) pre-treatment (B). Cell index was recorded every 15 minutes (n=4 independent isolations, in duplicate) and statistical analyses were performed by functional ANOVA (R package fdANOVA). (C and D) Mitochondrial biogenesis was quantified in NCMs controls (DMSO + PBS) treated with or without MitoQ (C) or EUK 134 (D) pre-treatment by RT-qPCR of PGC1α, NRF 1 and NRF2. Data were normalized to HPRT. (E) Mitophagy was quantified in NCMs controls (DMSO + PBS) treated with MitoQ pre-treatment by western blot of parkin, ubiquitinylated proteins, beclin-1, LC3I, LC3 II and LC3II / I ratio. Data were normalized to actin. Images of western blot in Figure 4E. (F) Autophagy was quantified in NCMs treated by Iso with or without MitoQ pre-treatment and with or without Bafilomycin A1 (Baf) co-treatment by western blot of LC3I, LC3 II and LC3II / I ratio. (G) Mitophagy was quantified in NCMs treated with or without EUK 134 pre-treatment by western blot of parkin, ubiquitinylated proteins, beclin-1, LC3I, LC3 II and LC3II / I ratio. Data were normalized to actin. *P* values are indicated with at least 3 independent experiments. Representative images were selected to represent the mean values of each condition.

**Figure S4:**
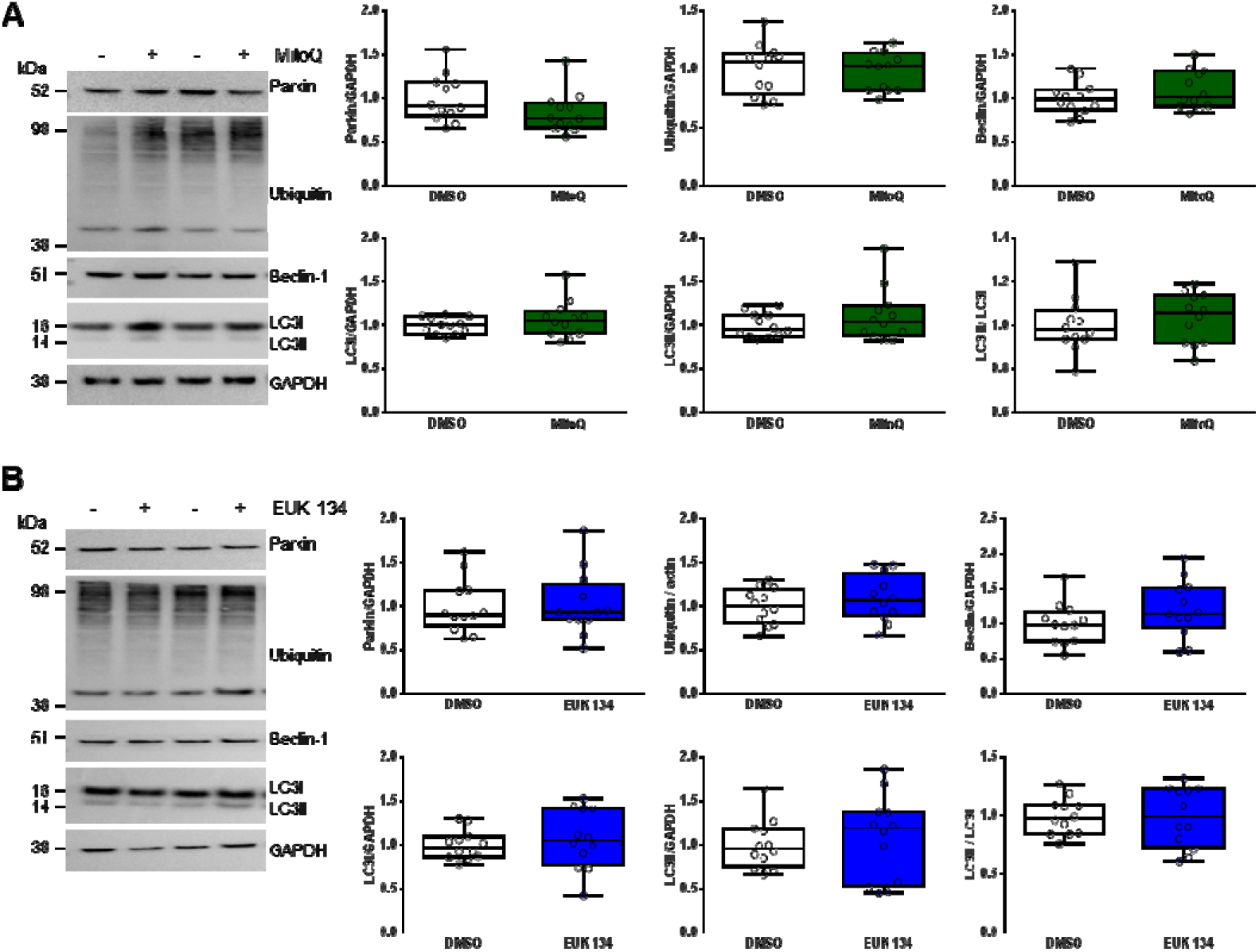
Effect of SOD mimic anti-oxydant (EUK 134) on mitophagy in neonatal rat fibroblasts. Mitophagy was quantified in NCFs treated with or without MitoQ (A) or EUK 134 (B) pre-treatments by western blot of parkin, ubiquitinylated proteins, beclin-1, LC3I, LC3 II and LC3II / I ratio. Data were normalized to GAPDH. Representative images were selected to represent the mean values of each condition.

